# Novel insights into the fundamentals of palatal shelf elevation dynamics in normal mouse embryos

**DOI:** 10.1101/2022.06.02.494562

**Authors:** Jeremy P. Goering, Michael Moedritzer, Marta Stetsiv, Dona Greta Isai, Brittany M. Hufft-Martinez, Zaid Umar, Madison K. Rickabaugh, Paul Keselman, Munish Chauhan, William M. Brooks, Kenneth J. Fischer, Andras Czirok, Irfan Saadi

## Abstract

Embryonic palate development involves bilateral vertical growth of palatal shelves – extensions from the maxillary processes – next to the tongue until embryonic day (E) 13.5. Following vertical growth, palatal shelves elevate and adhere above the tongue by E14.5. Current models indicate that this process of elevation involves a complex vertical to horizontal reorienting of the palatal shelves. While earlier studies have implied that this is a rapid process, the precise timing has not been resolved. To understand the dynamics of palatal shelf elevation, we employed time-restricted pregnancies with a one-hour resolution and magnetic resonance imaging of intermediate stages. Our data showed that in almost all C57BL/6J embryos, palatal shelves have not yet elevated by E14.0. However, six hours later at E14.25, palatal shelves have completed elevation in 80% of embryos. Interestingly, all E14.25 embryos with unelevated palatal shelves (20%) were female, suggesting a delay in female embryos. In FVB/NJ embryos, the elevation window started earlier (E13.875-E14.25) without any noticeable sex differences. We frequently captured an intermediate stage with unilateral elevation of either right or left palatal shelf. Magnetic resonance imaging of various stages showed that palatal shelf elevation began with the formation of bilateral bulges in the posterior. These bulges progressed laterally and anteriorly over time. During elevation, we observed increased cell proliferation in the lingual region of the palatal shelf. Within the bulge, cell orientation was acutely tilted towards the tongue and actomyosin activity was increased, which together may participate in the projection of the bulge in the horizontal direction. Thus, our data reveal novel insights into the rapid dynamic changes in palatal shelf elevation that lay the foundation for future studies of normal and abnormal palatogenesis.

## Introduction

Clefts of the lip and palate constitute the majority of craniofacial malformations, which collectively affect ~1/800 births worldwide (Mossey et al. 2009; Mossey and Modell 2012). Clefts of the palate alone affect ~1/1700 of all births and are more common among females than males (Mossey et al. 2003; Mossey et al. 2009). Approximately half of these cleft palate only cases are isolated or nonsyndromic occurrences that have a complex etiology, with both genetic and environmental factors (Dixon et al. 2011; Jugessur et al. 2009; Leslie et al. 2016; Leslie et al. 2015; Mangold et al. 2011; Martinelli et al. 2020). The contribution of these factors has been extensively studied using rodent models over the past six decades. The initial focus of these studies was on environmental factors and chemical compounds as well as on susceptible murine backgrounds and spontaneous mouse mutants (Greene and Kochhar 1973; 1975; Juriloff 1980; Pratt et al. 1984a; Pratt et al. 1984b; Slavkin and Melnick 1982; Trasler 1960; Vekemans and Biddle 1984).

Another early focus was to determine normal palatogenesis. Classical studies identified three main steps in palatogenesis following palatal shelf (PS) induction: 1) vertical PS growth next to the tongue, 2) PS elevation above the tongue, and 3) PS fusion in the midline (Diewert and Wang 1992; Ferguson 1988; Gritli-Linde 2007; Johnston et al. 1975; Walker and Fraser 1956b). These steps were confirmed in humans through fetal analysis as well as improved ultrasound imaging and are thus part of most textbooks (Burdi 1965; Burdi and Silvey 1969; Diewert 1983; 1985; Diewert and Shiota 1990).

Among these three steps, PS elevation has remained enigmatic. Early studies of PS elevation suggested a simple rotation of the vertical PS to a horizontal position, followed by horizontal growth with cell proliferation. This model was challenged by Walker and Fraser (1956) (Walker and Fraser 1956a) who showed that the process was too rapid – less than two minutes with manual manipulation – for cell proliferation to be the main driver of elevation. They further argued that the PS moved from vertical to horizontal direction via a process of reorientation rather than simple rotation. They also proposed that palatal shelf elevation occurred in a developmental window approximately 3 hours long, but noted that this developmental window was shifted in different mouse strains.

Since these early studies, there were few attempts to study normal PS elevation, though some insights were gained from the few mouse mutants that appeared to affect PS elevation. The current understanding of PS elevation was comprehensively summarized by Bush and Jiang (2012) (Bush and Jiang 2012). According to the model they compiled, the anterior-most regions of the PS elevated by rotating, while the middle and posterior PS regions elevated by reorienting from a vertical to a horizontal position, as also proposed by Walker and Fraser (1956). These determinations were based on location of the medial edge epithelium (MEE), which eventually meets and fuses in the midline in the elevated shelf. Prior to elevation, the MEE in the anterior-most region of the PS lies at the ventral tip of each vertical shelf. In contrast, the MEE in the middle and posterior PS regions lies on the lateral surface. Several pieces of evidence suggest that the vertical to horizontal reorienting initiates with a bulge formation that extended above the tongue. Bush and Jiang (2012) also noted Walker and Fraser’s assertion that this remodeling may proceed one PS at a time, as unilateral PS elevation was occasionally observed in normal mouse embryos.

We recently reported that deficiency of *Specc1l* resulted in PS elevation delay, but the PS shelves still eventually elevated and fused (Goering et al. 2021b; Hall et al. 2020). The PS elevation delay provides a model for nonsyndromic cleft palate where a delay in elevation can be considered a sensitized background that combined with additional negative genetic or environmental factors, may put the affected individual above the threshold for isolated cleft palate.

While studying the delay in PS elevation in our *Specc1l* mutant alleles, we encountered a lot of inter-litter variability in timing and staging of the shelves. To better understand normal PS elevation, we performed a careful analysis of two commonly used murine strains (C57BL/6J and FVB/NJ) with a 3-hour resolution of embryonic development using time-restricted matings. Our data show that: 1) the PS could elevate in <3 hours, 2) there are mouse strain and sex differences in the embryonic PS elevation, and 3) vertical to horizontal remodeling occurs with dynamic lateral anteroposterior changes in the PS. These results provide critical insights that will not only help understand normal PS elevation, but also help characterize PS elevation defects in existing and novel mutant mouse models.

## Results

### Palatal shelves elevate between E14.0-E14.25 in C57BL/6J embryos

The most important aspect in determining the PS elevation dynamics was to control the inter-litter variability in embryonic age. We decided to focus on the timing of conception as a source of this variability. Compared to standard overnight matings, we performed time-restricted matings where we checked vaginal plugs every hour. All data presented here are from such time-restricted matings with a 1-hour resolution. We began by identifying the latest embryonic timepoint at which the PS were still vertical or unelevated in C57BL/6J embryos. The convention in the field is to consider embryonic day (E) 13.5 from overnight matings for unelevated PS and E14.5 for elevated PS, which represents a 24-hour period. We decided to score embryos from various time-restricted matings into three categories or stages of PS elevation: bilaterally unelevated **(Fig.1A)**, unilaterally elevated **(Fig.1B)**, and bilaterally elevated or adhering PS **(Fig.1C)**. We found that at E14.0, almost all (97%, n=38) PS were still unelevated **(Fig.1D, E14.0)**. We then looked backwards from E14.5 for the earliest timepoint where most PS were elevated. We found that at E14.25, 80% (n=64) of embryos showed bilateral PS elevation, while another 11% showed a unilateral PS elevation **(Fig.1D, E14.25)**. Thus, our results indicated that in C57BL/6J embryos PS elevation is largely completed in less than 6 hours.

**Figure 1:**
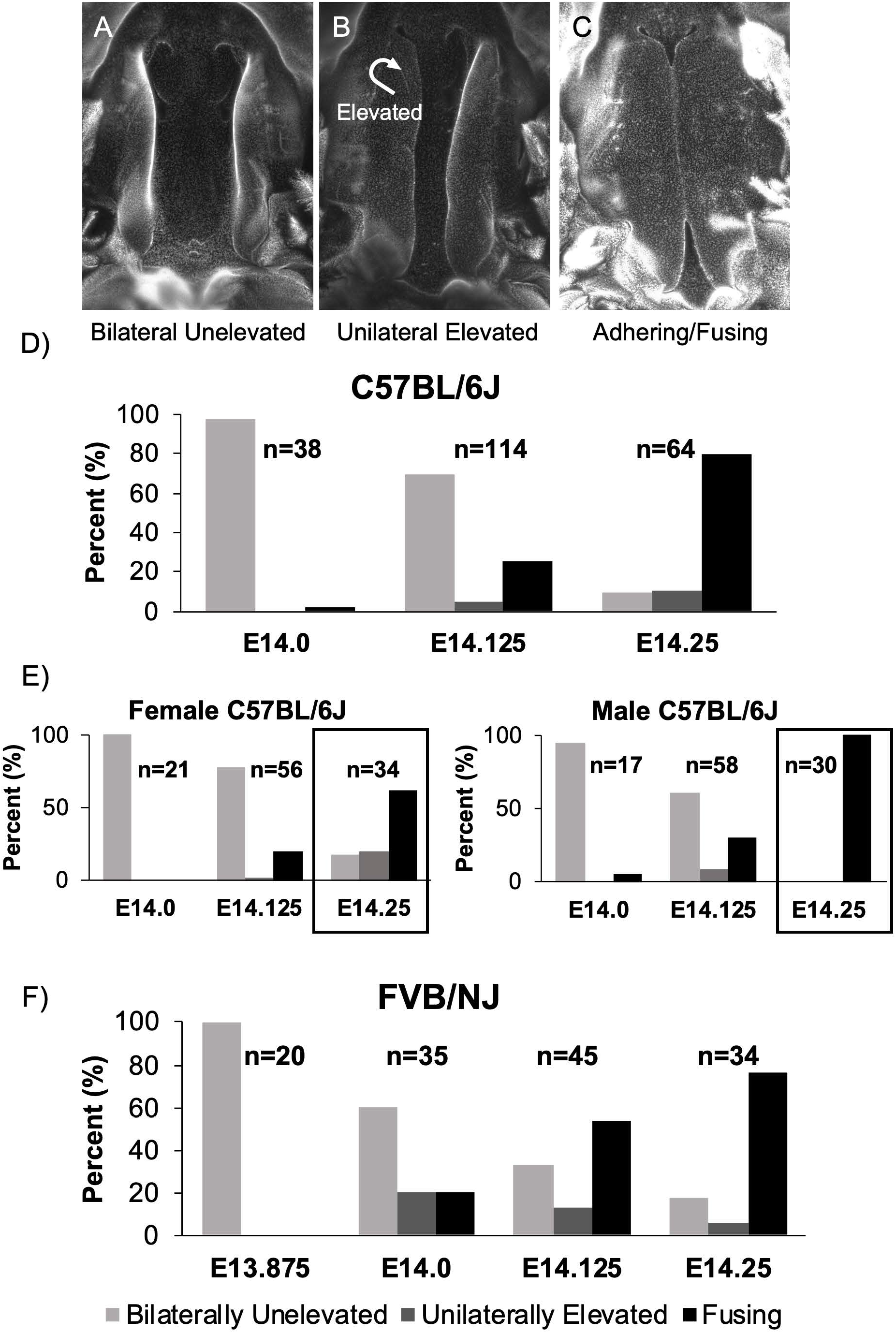
Time-restricted matings show that palatal shelves elevate in less than 6 hours. **A-C)** To carefully time palatal shelf closure in C57BL/6J and FVB/NJ embryos, we used time-restricted matings where we checked plugs every hour to reduce inter-litter variability in embryonic development. We decided to score the palatal shelves for 3 stages: Bilaterally unelevated (A), Unilaterally elevated (B) and Bilaterally elevated or Adhering (C). **D)** We found that at E14.0 in C57BL/6J embryos, 97% (37/38) of palatal shelves were bilaterally unelevated. Next, we found that at E14.25, just 6 hours later, 80% (51/64) of embryos showed bilaterally elevated palatal shelves. An additional 11% of embryos at E14.25 showed unilaterally elevated palatal shelves. Finally, we found that even at E14.125, 21% of embryos had completed palatal shelf elevation, indicating that once initiated palatal shelves could complete elevation in less than 3 hours. **E)** We also determined the sex of the embryos we isolated. We found that at E14.25, 100% (30/30) male embryos showed bilaterally elevated palatal shelves. In contrast, 18% (6/34) of female embryos at E14.25 showed bilaterally unelevated palatal shelves (boxed regions). In fact, all C57BL/6J embryos with bilaterally or unilaterally unelevated palatal shelves at E14.25 were female (D vs. E). **F)** In comparison to C57BL/6J, the FVB/NJ embryos showed a broader window for palatal shelf elevation. At 14.0, 20% (7/35) of FVB/NJ embryos had bilaterally elevated palatal shelves and another 20% showed unilaterally elevated palatal shelves. However, at E14.25 76% (26/34) of FVB/NJ embryos showed bilaterally elevated palatal shelves, similar to the 80% observed in C57BL/6J embryos.

To further assess the process of PS elevation, we looked at embryos at the intermediate E14.125 timepoint **(Fig.1D, E14.125)**. We found that 25% of embryos (n=114) had already completed PS elevation. Thus implying that once initiated, PS elevation needed <3 hours to complete, as previously suggested by Walker and Fraser (1956) (Walker and Fraser 1956b). Interestingly, we only observed 5% of embryos with unilateral PS elevation at E14.125, suggesting that it is not a stable intermediate state. Furthermore, either right or left PS was equally observed to be unilaterally elevated, indicating a random process. Overall, our data depict PS elevation as a rapid embryonic process, which may or may not have an obligatory intermediate unilateral elevation state.

### Female C57BL/6J embryos showed a delay in PS elevation

We next wanted to determine if there were any sex differences in this PS elevation process in C57BL/6J embryos. At E14.25, 20% of embryos had not yet completed PS elevation, including 9% with bilateral unelevated PS and 11% with unilateral elevated PS **(Fig.1D, E14.25)**. Interestingly, all these embryos with incomplete PS elevation were female, constituting 39% of female embryos at E14.25 (n=34) **(Fig.1E, Female, E14.25)**. In contrast, 100% of male embryos (n=30) had completed PS elevation at E14.25 **(Fig.1E, Male, E14.25)**. Even at E14.125, 78% of female embryos had not initiated PS elevation (n=56) compared with 61% of male embryos (n=58). Thus, our data suggest that a significant proportion of female C57BL/6J embryos have delayed PS elevation.

### Comparison of palate elevation dynamics in FVB/NJ embryos

Strain differences in palate closure dynamics have been reported previously. These studies usually compared C57BL/6 strain to cleft palate susceptible strains such as A/J and S/Wyn. These susceptible strains invariably showed delayed PS elevation (Ciriani and Diewert 1986; Diewert 1982; 1986; Fraser 1976; Walker and Fraser 1956b). We wanted to compare another common laboratory strain, FVB/NJ, that is not known to be susceptible to cleft palate. We found two main strain differences. First, the PS elevation started earlier in FVB/NJ compared to C57BL/6J embryos. At E14.0, 40% of FVB/NJ embryos (n=35) already had bilaterally or unilaterally elevated PS **(Fig.1F, E14.0)**. We confirmed that three hours earlier, at E13.875, 100% of FVB/NJ embryos (n=20) had unelevated PS **(Fig.1F, E13.875)**. Second, we did not observe sex differences in FVB/NJ embryos at E14.25 **(Fig.1F, E14.25)**. Female FVB/NJ embryos did not show a delay in PS elevation compared to males or to overall C57BL/6J data **(Fig.S1)**. At E14.25, the overall extent of PS elevation was similar between C57BL/6J and FVB/NJ embryos **(Fig.1D,F)**.

### Anteroposterior dynamics of PS remodeling suggest a posterior to anterior elevation

Previous studies have highlighted anteroposterior differences in PS elevation (Bush and Jiang 2012; Chiquet et al. 2016; Liu et al. 2021; Walker and Fraser 1956b; Yu and Ornitz 2011). Given our ability to consistently obtain embryos in the process of PS elevation, we were able to collect and scan several of them using magnetic resonance imaging (MRI). These scans were segmented and 3D images generated to get a better anteroposterior view of the PS during elevation. The 3D images showed that PS elevation does progress through lateral bulges above the tongue **(Fig.2)**. However, these lateral bulges were first observed and most prominent in the posterior palate **(Fig.2A, arrows)**. We also observed that the bulges gradually tapered anteriorly **(Fig.2B, arrowheads)**. In this example **(Fig.2B)**, we observed the bulges to be similarly extended bilaterally with yet limited extension in the mid-palate region. We noticed that most instances of lateral bulges at mid-palate region **(Fig.2C, arrowheads; Fig.S2)** were in embryos with unilaterally elevated PS **(Fig.2C, arrow)**. Interestingly, the very posterior ends of the PS in embryos with unilateral elevation all appeared to be already bilaterally horizontal. Nonetheless, in all instances captured, we observed that PS adhesion began at anterior to mid-palate region and extended posteriorly **(Fig.2D; Fig.S2)** – consistent with current understanding. Taken together, these results showed that PS elevation – in contrast to adhesion – proceeded in the posterior to anterior direction.

**Figure 2:**
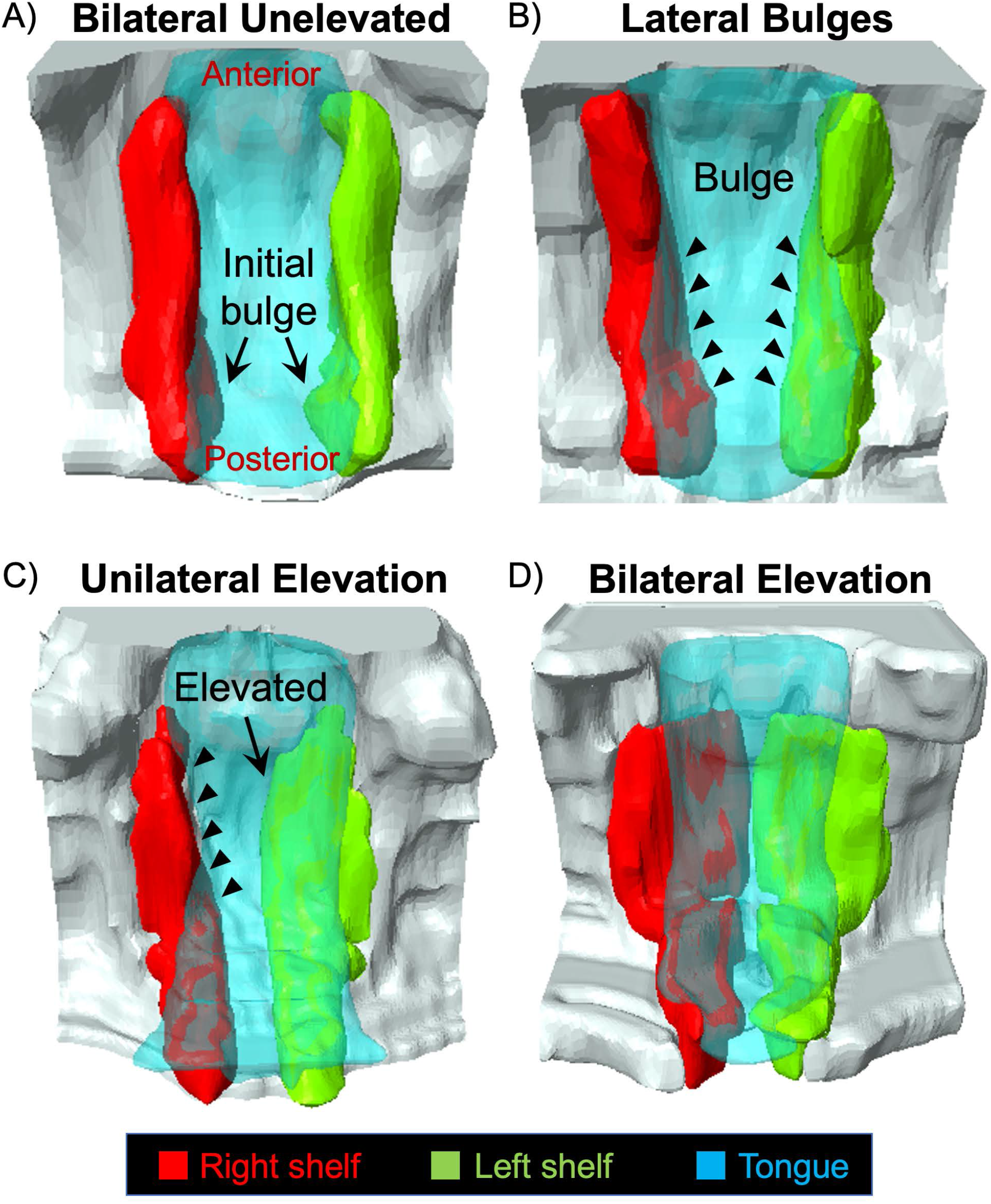
3-D Imaging reveals anteroposterior dynamics of palatal shelf elevation. We used magnetic resonance imaging (MRI) scans of the embryos isolated at various stages of palatal shelf elevation to segment and construct 3-D images. Among the images of largely bilaterally unelevated palatal shelves, we noticed lateral bulges appearing bilaterally in the very posterior part of the palatal shelves (A, arrows). In other scans we observed more extensive lateral bulges (B, arrowheads). Again, these bulges were most prominent in the posterior palate and gradually tapered anteriorly (B, arrowheads). We also scanned unilaterally elevated palatal shelves (C, arrow). The unelevated palatal shelf is these instances showed the most prominent lateral bulge extending into the middle and anterior palate (B vs. C, arrowheads). Panels A-C clearly showed that the lateral bulges progressed from the posterior to anterior direction. **D)** Scans of bilaterally elevated palatal shelves showed that adhesion first occurred in the anterior palate and proceeded posteriorly as expected.

### Regional changes in cell proliferation observed during PS elevation

Once we could identify and capture the PS bulge region consistently, we looked at cell proliferation in the bulged PS. We analyzed the PS entirely as well as in regions – lingual, buccal and hinge – where the bulge develops in the hinge region **(Fig.3)**. We observed increased cell proliferation only in the lingual region, but this increase was present prior to elevation in the unelevated PS and persisted till the elevated PS **(Fig.3A, graph)**. This result suggested that vertical to horizontal remodeling during PS elevation does not require cell proliferation within the bulge itself, but that increased cell proliferation in the lingual region may facilitate the process.

**Figure 3:**
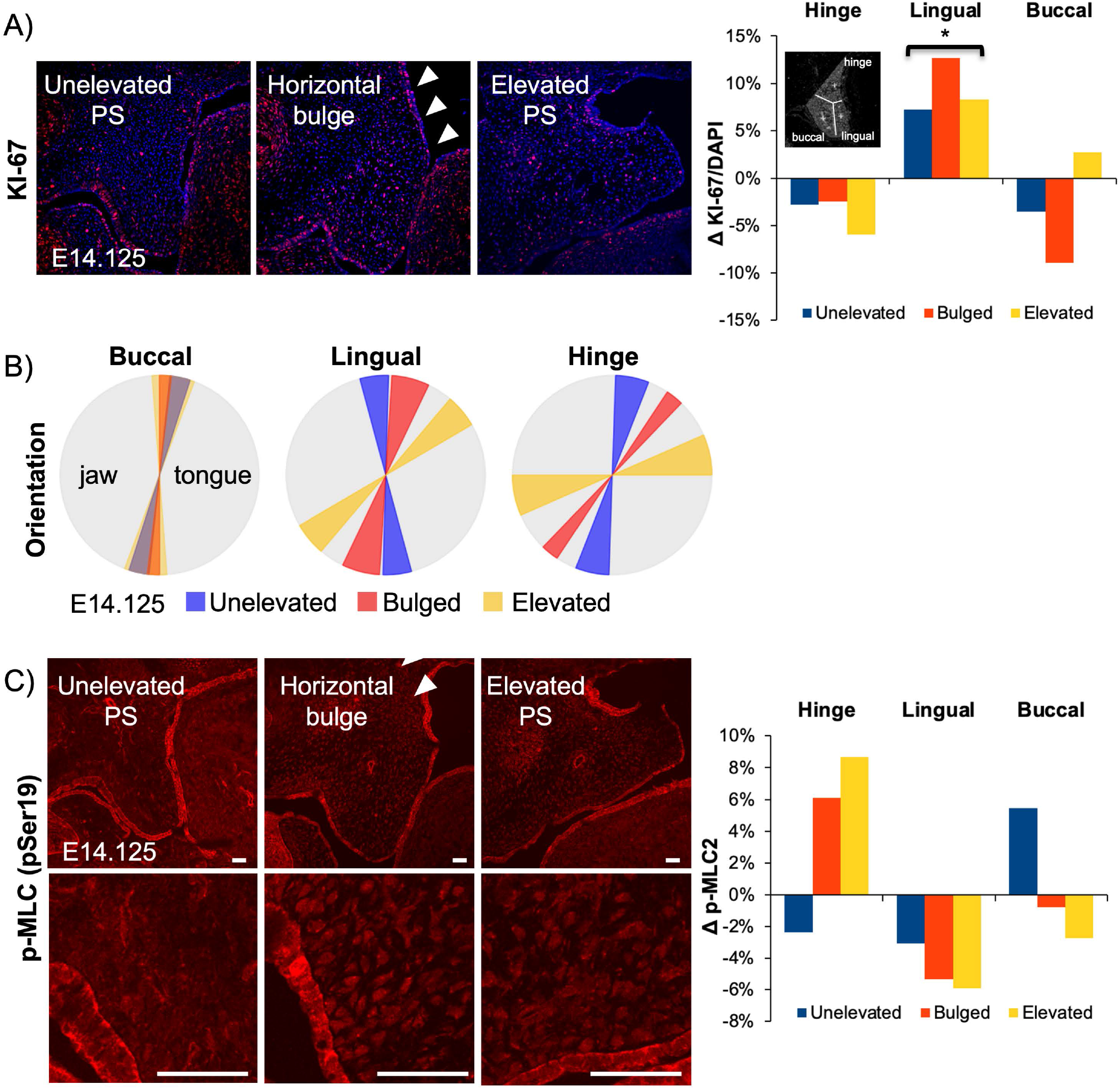
Changes in mesenchymal cell proliferation, orientation and actomyosin contraction during palatal shelf elevation. Cellular changes during palatal shelf elevation were characterized by assessing mesenchymal cell proliferation **(A)**, cell orientation **(B)** and phospho-Myosin Light Chain 2 (p-MLC2) level **(C)**. **A)** Cell proliferation was assessed with anti-KI-67 antibody staining in cryosections from E14.125 embryos with unelevated, bulged and elevated palatal shelves. Entire palatal shelves were analyzed to determine the mean. Differences from the mean were then plotted for three regions of the palatal shelves: hinge, lingual and buccal (inset in graph). Only the lingual region showed a statistically significant (p<0.05) increase in cell proliferation throughout unelevated, bulged and elevated palatal shelves. **B)** Cell orientation was assessed by determining the nuclear shape and angle. Local and regional averages and standard deviation were plotted as a wedge diagram. A tilt to the right or left signifies orientation towards the tongue or the jaw, respectively. The three regions of the palatal shelf (buccal, lingual, hinge) were analyzed separately. Cells in the buccal region showed a slight tilt towards the tongue, which stayed similar throughout palate elevation. Thus, cells in the buccal region do not participate in the vertical to horizontal remodeling. Cells in the lingual region showed a largely vertical orientation in an unelevated shelf, a slight tilt towards the tongue in a bulged palatal shelf, and a large 60° tilt in the elevated shelf, indicating a significant participation in the horizontal remodeling. Cells in the hinge region showed the most aggressive tilt towards the tongue during the three stages, with an almost horizontal orientation in the elevated palatal shelf. **C)** Activated p-MLC2 staining was assessed as a proxy for actomyosin activity. Qualitative changes appear to show more condensed expression in the bulged and elevated palatal shelves (magnified images, lower panels). Quantitative analysis was assessed as a change from average expression for each region. Spatially increased p-MLC2 expression was first observed in the buccal region in the unelevated shelf. Later, in bulged and elevated palatal shelves, the increased activity was observed in the hinge region. The lingual region showed consistently lower than average activity throughout palate elevation.

### Cell orientation changed drastically in the lateral bulge during PS elevation

We previously showed that prior to PS elevation at E13.5, the PS mesenchymal cells are aligned and slightly oriented towards the tongue (lingual direction) (Goering et al. 2021a). Thus, we wanted to assess changes in cell orientation during PS elevation **(Fig.3B)**. We observed that the only significant changes in cell orientation occurred in the hinge region **(Fig.3B)**. The cells in the hinge region were already more oriented towards the tongue in bilaterally unelevated PS. In the bulged PS, the mesenchymal cells were acutely oriented towards the tongue **(Fig.3B, Hinge, red)**. As expected, in the elevated PS, the cells were almost horizontal both in the hinge and lingual regions **(Fig.3B, yellow)**. Interestingly, the buccal region cells did not show any change in cell orientation, indicating that these cells are largely excluded from the horizontal or elevated part of the PS **(Fig.3B, Buccal)**.

### Sequential increase in activated myosin light chain levels in buccal and hinge regions during PS elevation

Next, we considered the possibility of increased actomyosin activity in the PS bulge region during PS elevation. We looked at levels of phosphorylated myosin light chain (p-MLC) that participates in both muscle and non-muscle myosin-based contractility. We observed evidence for a bipartite change in p-MLC levels **(Fig.4C)**. In the unelevated PS, we observed an increase in the buccal region **(Fig.4C, graph, buccal)**. However, later we observed an increase within the bulge in the hinge region **(Fig.4C, graph, hinge)**. The increased activity in the hinge region persisted in the elevated PS. This observed 2-step pattern is consistent with an initial vertical PS contraction and a subsequent horizontal contraction, which together may propel the bulge in the horizontal direction.

**Figure 4:**
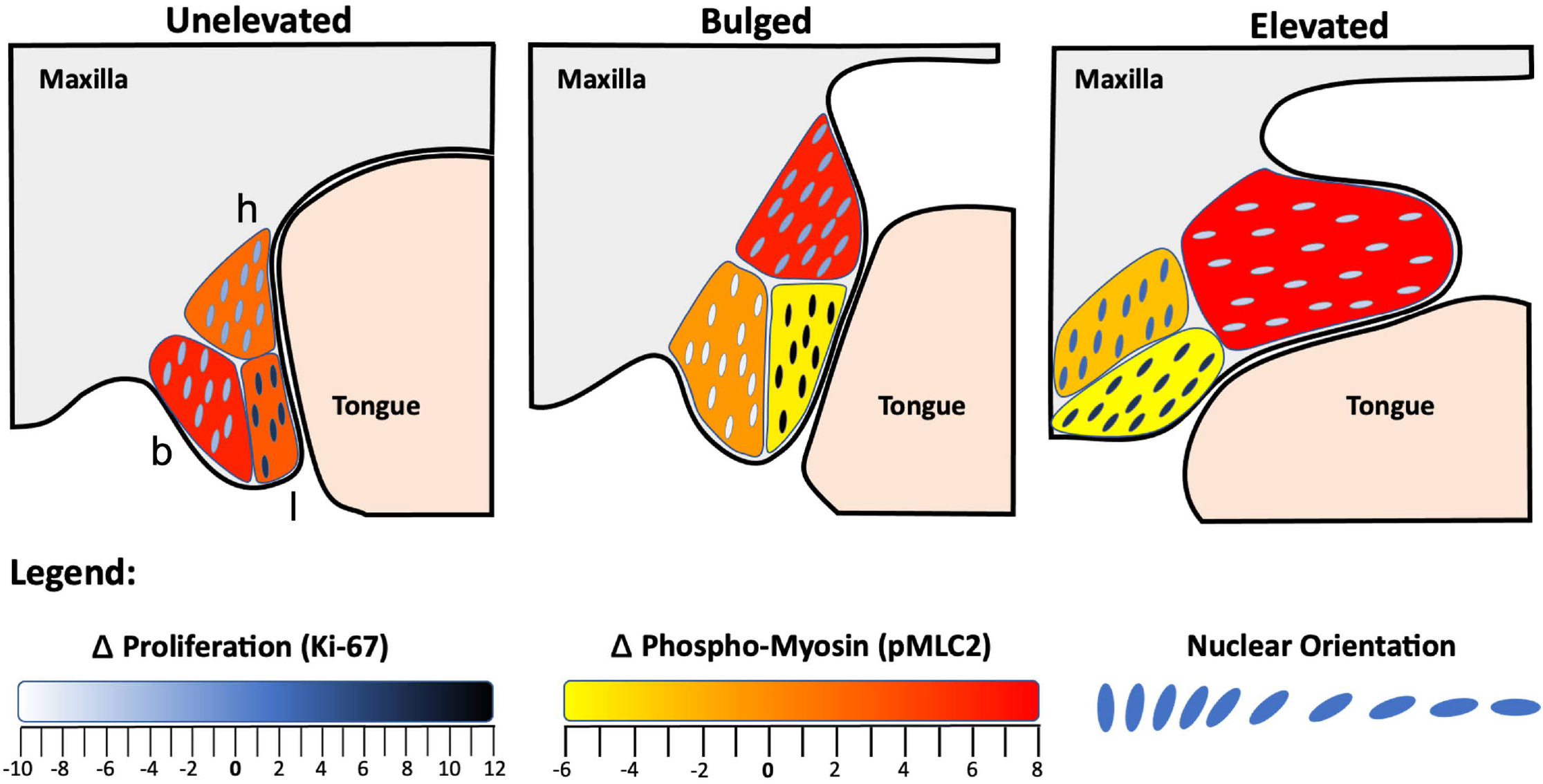
Model of palatal shelf elevation. Schematic summarizing the cellular changes observed in the buccal (b), lingual (l) and hinge (h) regions of the unelevated, bulged and elevated palatal shelves. Immediately prior to elevation, we observed increased cell proliferation in lingual region, a slight tilt in orientation towards the tongue in both buccal and hinge regions, and increased actomyosin activity in the buccal region. The latter may participate in vertical contraction of the palatal shelf. In the bulged palatal shelf, increased cell proliferation persisted in the lingual region, cell orientation was acutely tilted towards the tongue within the bulge in the hinge region, and actomyosin activity was now increased in the hinge region. The acute orientation and increased actomyosin activity in the hinge region may participate in the propulsion of the bulge in the horizontal direction. In the elevated palatal shelf, cell proliferation continued in the posterior lingual region, the cell orientation in the lingual and hinge regions are almost horizontal, and actomyosin activity is still increased in the hinge region. The lack of change in the cell orientation of buccal region suggests that cells in that region did not participate in the vertical to horizontal remodeling. The spatial pattern of actomyosin activity suggested that vertical palatal shelf contraction preceded in the buccal followed by dorsal contraction in the hinge region which coincided with the horizontal bulge formation. It remains to be seen whether the ventral portion of the hinge region and proliferating lingual cells are the ones that are propelled into the elevated palatal shelf.

## Discussion

Even though palatogenesis has been studied extensively, the actual timing and sequence of events of PS elevation have remained relatively elusive. While the convention in the field is that PS elevation occurs between E13.5 and E14.5, several studies have shown that it is a more rapid process. Perhaps the most elegant and earliest of these studies were by Walker and Fraser (1956) (Walker and Fraser 1956b) who proposed that PS elevation *in utero* may take place in ~3 hours. Using more precise timed-matings with reduced inter-litter variability, we now show that once initiated PS elevation can complete in <3 hours *in utero*. In C57BL/6J embryos, our data indicated a developmental window of E14.0-E14.25 (6 hours) wherein almost all PS elevation is completed.

Strain differences have previously been studied in palatogenesis, particularly for strains that showed increased occurrence of cleft palate. In general, these studies showed a delay in palate elevation in strains susceptible to cleft palate, e.g. A/J and A/WySnJ, compared to C57BL/6J (Ciriani and Diewert 1986; Diewert 1982; 1986; Fraser 1976; Walker and Fraser 1956b). We compared the FVB/NJ strain, which is not reported to have increased susceptibility to cleft palate. We found that the overall window of PS elevation is broader in FVB/NJ (~9 hours; E13.875-E14.25) compared to C57BL/6J (~6 hours; E14.0-E14.25) embryos, however, at E14.25, a similar number of embryos had completed PS elevation in both strains.

A surprising finding in our results was the identification of sex differences in the timing of PS elevation. Cleft palate only in humans is known to occur more frequently in females vs males (~2:1) (Mossey et al. 2003; Mossey et al. 2009). And there have been early studies suggesting delayed PS elevation in human (Burdi and Silvey 1969) and mouse (Burdi and Faist 1967) female embryos. We found that almost 39% of C57BL/6J female embryos had not completed PS elevation by E14.25, whereas 100% of C57BL/6J male embryos showed completed PS elevation. We also recently reported a mouse model (*Specc1l^ΔCCD2^*) on a mostly C57BL/6J background (N3 generation) mixed with some FVB/NJ that the cleft palate phenotype in heterozygotes was more prevalent in female (17%) vs. male (5%) embryos (Goering et al. 2021b). In fact, we saw a drastic decrease in *Specc1l^ΔCCD2/+^* heterozygous females as we further backcrossed to C57BL/6J background, with almost no *Specc1l^CCD2/+^* females at weaning in the N7 generation. These *Specc1l^ΔCCD2/+^* results argued that the C57BL/6J background predisposed females for cleft palate. Our data in this paper showing a delay in C57BL/6J female embryos are consistent with the above argument. However, we want to emphasize that we are not proposing that the FVB/NJ background offers any protection against cleft palate. In fact, our preliminary data suggest a similar overall incidence of cleft palate in *Specc1l^ΔCCD2/+^* heterozygotes (~20%) on a pure FVB/NJ background (>N8 generation), however, with equal occurrence in male or female embryos (not shown). Thus, sex differences in PS elevation represent one of many factors that affect palatogenesis and need to be considered carefully in future studies.

Our data clearly showed that the PS undergo vertical to horizontal remodeling via lateral bulge formation. However, our imaging of intermediate states indicated that the bulges originated in the posterior palate and gradually extended anteriorly. In contrast, PS adhesion eventually began in the anterior to mid-palate region and extended posteriorly, exactly as expected. Our finding is consistent with the Walker and Fraser (1956) proposal that the elevation force that creates the lateral bulges is generated from the posterior region of the palate, and that the greater thickness of the tongue in the posterior slows PS elevation.

Another intriguing speculation made by Walker and Fraser (1956) was that the unilateral PS elevation is an “obligate” intermediate step. This would imply that one PS elevates above the tongue prior to the other shelf and that the tongue goes through a rocker like motion. There have been a few subsequent studies where unilaterally elevated PS have been captured in normal embryos (Bush and Jiang 2012; Liu et al. 2021; Yu and Ornitz 2011), and several studies where unilateral elevation was captured in mutant mouse models of cleft palate, including those with *Specc1l* deficiency (Butali et al. 2019; Goodwin et al. 2020; Hall et al. 2020; Hill et al. 2015; Liu et al. 2008). We were keen to see if we would routinely capture unilaterally elevated PS in our study. While we did capture many instances of unilateral PS elevation, we also encountered several with equally bilateral bulges. However, when compared to the instances where the shelves were equally bilaterally elevated, the unelevated PS in embryos with unilateral PS elevation had a more anteriorly progressed bulge. Thus, we argue that the bulges initially developed bilaterally and when they progressed to a certain extent, unilateral elevation took place. We did not find any preference for right or left PS elevation among unilaterally elevated samples. Thus, if unilateral elevation is indeed an “obligate” intermediate step, it is very transient and happens late in the elevation process. A normal process that proceeded unilaterally would be more consistent with the human condition as well, as occurrence of unilateral human clefts is common. Further, a mechanical force generated from the posterior regions of the palate may not be evenly split and a one-at-a-time PS elevation may allow for more flexibility and reduced force requirement to displace the tongue.

Lastly, the nature of PS elevation has long been debated (Bush and Jiang 2012; Ferguson 1977; 1988; Lazzaro 1940; Walker and Fraser 1956b). The vertical to horizontal remodeling was thought to involve rotation as well as cell proliferation. While Walker and Fraser (1956) opposed both, it remained to be seen if cell proliferation played a role. We now show that the PS bulges during elevation (hinge region) do not indicate any relative increase in cell proliferation. However, we did observe increased cell proliferation in the lingual region, which was already present in the unelevated PS and persisted through to the elevated PS. Thus, this increased cell proliferation in the lingual region, which is immediately ventral to the hinge region where the bulges form, may passively participate in the PS elevation. It also suggests that there is no horizontal growth following elevation. Instead, the vertical growth prior to PS elevation persists in the lingual region and culminates with PS elevation and adhesion.

Instead of increased cell proliferation in the PS bulge, we observed an increase in cell orientation towards the tongue and in activated myosin light chain staining. Thus, cell orientation and actomyosin forces may participate in the rapid movement of PS bulges during elevation. We previously showed that the PS mesenchymal cells were slightly oriented towards the tongue immediately prior to elevation at E13.5 (Goering et al. 2021a). We now show that the sharpest change in orientation towards the tongue occurs within the bulge in the hinge region. Following PS elevation, the lingual and hinge regions showed almost horizontal cell orientation, as expected, but the buccal region cells did not. This latter result suggested that the buccal region did not participate in the horizontal remodeling. Instead, we observed an increase in activated myosin light chain staining in the buccal region of the unelevated PS, suggesting that the buccal region may participate in the vertical contraction of the PS as the lateral bulges appeared. The combination of cells oriented towards the tongue, coordinated actomyosin contractility and permissive extracellular matrix conditions, allow for rapid reorienting of the shelves above the tongue. Future studies with mouse mutants with defects in PS elevation, cell orientation or alignment, and actomyosin contractility will help further delineate this process. We assert that PS elevation is the most dynamic and sensitive step in palatogenesis and a perfect target for both genetic and environmental factors in the etiology of cleft palate.

## Materials and Methods

### Time-restricted mouse matings and embryo processing

For time-restricted matings, a male and female mouse were placed in a cage together for one hour, then the female was visually checked for the presence of a vaginal plug. The time of plug observation was recorded, and if no plug is observed the female could be placed back with the male for one more hour. These hourly plug-checks provide a precise time-of-mating with a temporal resolution of one hour. The precise age of the embryos was determined relative to the time of the plug observation. At the desired embryonic time point, each pregnant female was euthanized using methods approved by IACUC. Time restricted matings were set-up in the morning. For example, if a plug was observed at 8:00AM, then the litter would be dissected 14 days later at 8:00AM for E14.0, or 11:00AM for an E14.125, or at 2:00PM for an E14.25. The embryos were harvested, washed in 1x PBS, then decapitated, jaw removed, and the palate elevation status was recorded. Yolk sacs were taken at the time of harvesting for genotyping. Sex was determined by PCR using primers flanking an 84 bp deletion of the *Rbm31x* gene relative to its gametolog *Rbm31y* (Forward: CACCTTAAGAACAAGCCAATACA; Reverse: GGCTTGTCCTGAAAACATTTGG). A single product (269 bp) was amplified in female subjects and two products (269 bp; 353 bp) amplified in male subjects (Tunster 2017).

### Whole-mount DAPI staining

Whole mount DAPI staining of the palate was achieved by decapitating fixed embryos, removing the lower jaw, and incubating the exposed palates in 500-1000nM DAPI solution overnight, then imaging on Nikon SMZ 1500 stereomicroscope (Goering et al. 2021a; Goering et al. 2021b; Sandell et al.).

### Static high-resolution magnetic resonance imaging (MRI) of embryos ex vivo

Embryos from time restricted matings were harvested at timepoints E14.0, E14.125, and E14.25 and fixed overnight in 4% PFA. The embryos were then incubated in 0.5mM MnCl2 for at least 24 hours. Embryos were then placed in 1x PBS in a 0.5mL microcentrifuge tube for scanning. Scanning was performed using a 10mm single loop surface RF coil and a 9.4 T Bruker (Avance Neo) system (Bruker, Billerica, MA, USA). High-resolution scans were acquired at a resolution between 20×25×170μm and 40×50×170μm.

### Segmentation and 3-D reconstruction

Image segmentation was performed using ScanIP (Synopsis) 3D analysis software. Masks for the palatal shelves and tongue were created by manually tracing the structures in each frame of the MRI slice-package. The masks were then used to generate 3D renderings in the ScanIP software.

### Immunofluorescence

Embryo heads were fixed in 4% PFA overnight, then submerged in 15% sucrose followed by 30% sucrose until the tissue sank. The head were then embedded in OCT media and coronal sections were obtained at a thickness of 10 μm. Antigen retrieval was performed by heating the slides in sodium citrate buffer (10mM Sodium Citrate, pH 6.0) at 96°C for 10 minutes. The slides were then washed in water and permeabilized with 0.5% Triton X-100 in 1x PBS for 30 minutes. After permeabilizing, the tissue was washed in 1x PBS for 3 x 5 minutes, then blocked in 10% normal goat serum (NGS) for 2 hours at room temperature. Tissue sections were incubated in primary antibodies KI-67 (CST, 12202) 1:500 and phospho-Myosin Light Chain Ser-1 (ECM Biosciences, MP3461) 1:100 overnight at 4°C then washed in PBS three times for 10 minutes each. Incubation in secondary antibody Goat anti-Mouse Alexa-Fluor 594 (Invitrogen, A11037) 1:500 was for two hours at room temperature. 0.1μM DAPI was incubated at the same time as secondary antibodies. 20x images were obtained on a Nikon Inverted Eclipse TiE with A1R confocal.

### Nuclear orientation assay

To quantitatively characterize local orientational ordering of mesenchyme nuclei, we performed the following image analysis sequence: 1) DAPI-labeled frozen sections were imaged with 20x objective magnification. 2) After loading the images into ImageJ (Schindelin et al.), the palatal shelf area was delineated, then nuclei were segmented by brightness-based global thresholding. 3) Touching nuclei in the binary segmented image were resolved by a watershed transformation. 4) Segmented clusters larger than 10 pixels were identified and fitted with an ellipse. 5) Nuclei with an unambiguous orientation (the ratio of the minor and major ellipse axes being less then 0.8) were assigned into cells of a spatial grid with a cell size of 50 um. 6) In each cell, we calculated the scalar 2D nematic order parameter (Das et al.) as *S*^2^ = 〈*cos2θ*〉^2^ + 〈*sin2θ*〉^2^, where theta (*θ*) is the angle between the long axis of the ellipse and a reference direction, and the averages 〈..〉 are calculated for each nucleus within the grid cell. The order parameter S=1 indicates a configuration in which nuclei are completely parallel, while S=0 indicates a completely random set. Intermediate numbers 0<S<1 indicate various degrees of partial ordering. Using the same data, we also calculated the direction of the local prevailing order. 7) We used the same measures to compare regions within the palatal shelves by pooling the segmented nuclei from multiple specimens into three spatial domains (lingual, buccal and hinge).

### Statistical analysis

To establish statistical significance, we calculated the quantitative measure for each independent sample (time-lapse recording, physical section of an embryo). The sets, containing at least 4 independent values, were compared by two-tailed Welch t-tests, which does not assume equal variance or paired data. For the calculations we used the scipy.stats.ttest_ind function of the python programming language.

## Acknowledgements

This project was supported in part by the National Institutes of Health grants DE026172 (I.S.), GM102801 (A.C.), and F31DE031181 (B.M.H.). I.S. was also supported in part by the Center of Biomedical Research Excellence (COBRE) grant (National Institute of General Medical Sciences P30 GM122731), Kansas IDeA Network for Biomedical Research Excellence grant (National Institute of General Medical Sciences P20 GM103418), and Kansas Intellectual and Developmental Disabilities Research Center (KIDDRC) grant (Eunice Kennedy Shriver National Institute of Child Health and Human Development, U54 HD090216). The Confocal Imaging Facility, the Integrated Imaging Core, and the Transgenic and Gene Targeting Institutional Facility at the University of Kansas Medical Center are supported, in part, by NIH/NIGMS COBRE grant P30 GM122731 and by NIH/NICHD KIDDRC grant U54 HD090216. The Leica STED microscope was supported by NIH S10 OD023625.

## Author contributions

JPG, MM, MS, WMB, PK, MC, KJF, AC, and IS conceived and designed the experiments. JPG, MM, MS, DGI, BMH, ZU, MKR, PK, and MC performed the experiments. JPG, PK, and MC performed the MR scans. DGI and AC performed the quantitative cell orientation and imaging analysis. KJF performed the FE analysis. JPG, KJF, AC and IS wrote the paper. MM, MS, BHM, MC and WMB edited the manuscript. All authors reviewed the manuscript.

