## Supplemental Figure 1 for "Novel insights into the fundamentals of palatal shelf elevation dynamics in normal mouse embryos"

Supplementary Figure 1:

A)

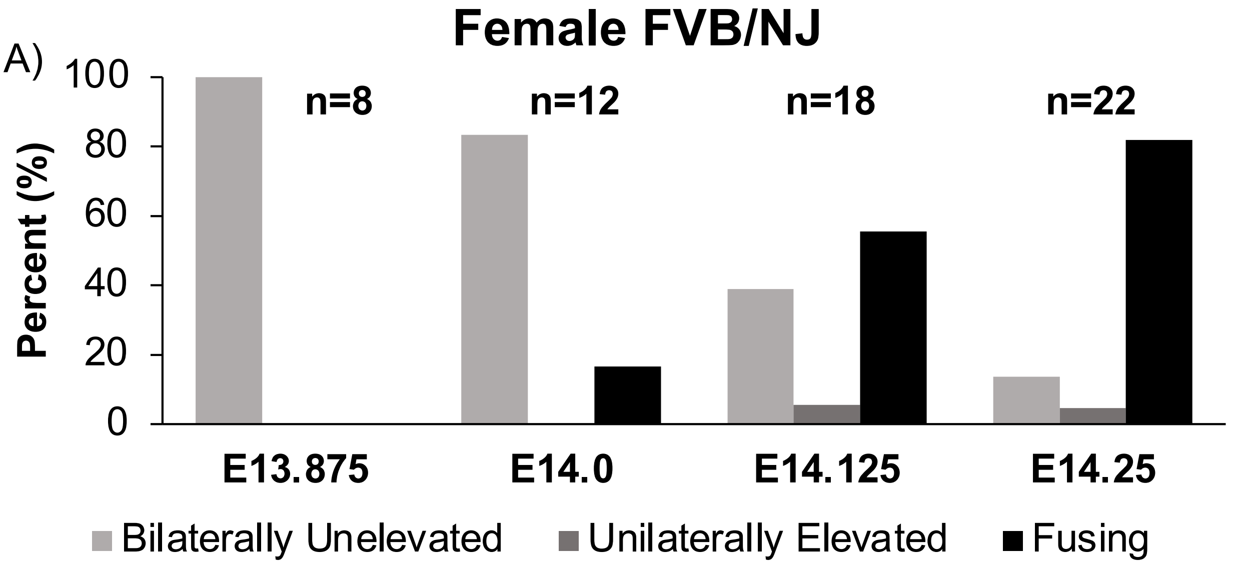


B)

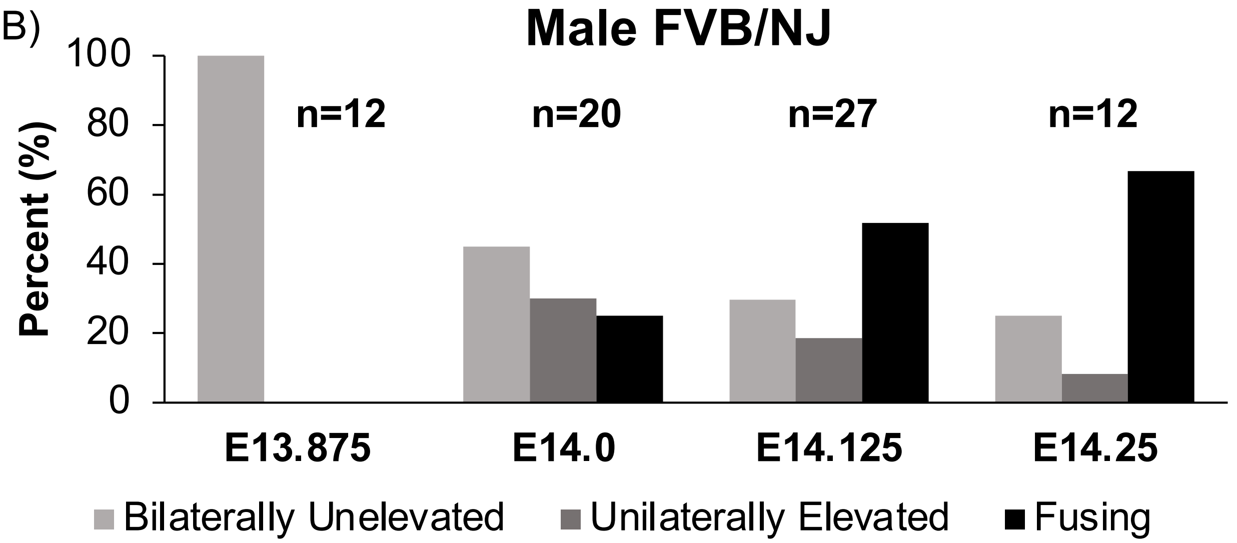


**Figure 1: FVB/NJ embryos did not show sex differences in the timing of palatal shelf elevation.** We determined the sex of the FVB/NJ embryos used in the data shown in **Figure 1F**. In contrast to C57BL/6J embryos, we did not find any significant sex differences among FVB/NJ embryos. At E14.0, there were ~80% female embryos that had unelevated palatal shelves (A, 14.0) compared to ~40% in males (B, 14.0). However, by E14.125, both male and female embryos showed ~50% elevated palatal shelves (A vs. B, E14.125). By E14.25, ~80% of female and ~70% of male embryos had elevated palatal shelves
